# Implicit agency is related to gamma power changes in an automatic imitation task

**DOI:** 10.1101/2021.06.28.448528

**Authors:** José Luis Ulloa, Roberta Vastano, Ole Jensen, Marcel Brass

## Abstract

Often we have a feeling that we can control effects in the external world through our actions. The role of action processing associated with this implicit form of agency is still not clear. In this study, we used automatic imitation and electroencephalography to investigate neural oscillations associated with action processing and its possible contribution to implicit agency. Brain activity was recorded while participants performed actions (congruent or incongruent with a displayed finger movement) which subsequently triggered an outcome (a tone). We used a time estimation task to measure intentional binding (an index of implicit agency). We observed a decrease of alpha, beta and gamma power for congruent compared to incongruent actions and increased theta power for incongruent compared to congruent actions. Crucially, participants who showed greater intentional binding for congruent versus incongruent actions also presented greater gamma power differences. Alpha, beta and theta power were modulated by congruency but were unrelated to intentional binding. Our study suggests that an increased implicit agency for facilitated actions is associated with changes in gamma power. Our study also contributes to a characterization of neural oscillations in automatic imitation.

## 1. INTRODUCTION

Agency is the experience of being in control of our actions and their consequences (Haggard and Tsakiris, 2009). Two aspects of agency have been investigated so far, an implicit “feeling of agency” and a more conscious “judgment of agency” (Synofzik et al., 2008, 2013). Implicit agency is related to the causal association between the actions we perform and the sensory consequences we perceive (Synofzik et al., 2008, 2013). This has been demonstrated in studies where voluntary (but not involuntary) actions and their sensory consequences are perceived occurring closer in time than when they are isolated (Haggard et al., 2002; Moore and Obhi, 2012). This perceived time compression (bringing action and consequence closer) is interpreted as the construction of a coherent conscious experience of our agency in time (Haggard, 2017; Haggard et al., 2002). This phenomenon is called intentional binding (IB) and it is said to reflect implicit agency. A predominant view states that agency emerges from mechanisms that compare intended actions and observed effects. However, a recent account suggests that mechanisms directly related to motor commands and the execution of actions might play a major role in agency (Chambon and Haggard, 2012; Chambon et al., 2014; Sidarus et al., 2017). These studies have shown that facilitating actions increases agency. In this vein, fluency of actions can be also modulated by imitative behaviour. In a study conducted by Vastano et al. (2017) agency was modulated with an automatic imitation task (Vastano et al., 2017, 2020). In this task participants had to lift their index or middle finger in response to an imperative cue while simultaneously observing a similar (congruent) or distinct (incongruent) finger movement of a mirrored right-hand (Brass et al., 2000, 2001; Cracco et al., 2018). Movement execution was thus facilitated by congruent and impeded by incongruent observed movements. Implicit agency was investigated measuring IB. Participants were asked to report the time elapsed between their actions and subsequent outcome tones. Intervals of time that were reported shorter relative to the real time intervals reflected a larger IB (Cravo et al., 2009; Kühn et al., 2013; Wen et al., 2015). Vastano et al. (2017) observed an effect of *congruency* on IB with more time compression for congruent relative to incongruent actions. In a follow-up EEG study these congruency effects correlated positively with late P300 changes and negatively with pre-response positivity changes, suggesting that modulations of cognitive load and interference impact implicit agency (Vastano et al. 2020).

Given the nature of automatic imitation it would be important to know if motor processes contribute to implicit agency in this task. There is a set of candidate neural oscillations associated with action processing that could be implicated in agency. Alpha band (8-12 Hz) oscillations generated in sensori-motor cortex are modulated by the execution and observation of movements (Cochin et al., 1999; Koelewijn et al., 2008; Muthukumaraswamy et al., 2004; Neuper et al., 2006; Pfurtscheller, 1981). Beta band (15–30 Hz) oscillations decrease rapidly several hundred milliseconds before an action and seem to reflect motor preparation (Cheyne et al., 2006; Engel and Fries, 2010; Gaetz et al., 2010; Wilson et al., 2010, 2014). Finally, gamma band (70–90 Hz) oscillations are seen around movement onset and seem to be associated with the action execution process itself (Ball et al., 2008; Cheyne et al., 2008; Crone et al., 1998a; Muthukumaraswamy, 2010; Pfurtscheller et al., 2003). Previous studies have shown a sensory and motor involvement of these neural oscillations in agency. For instance, participants that are exposed to ambiguous sensory feedback from their actions show changes in alpha oscillations in sensory areas (Kang et al., 2015) or changes in motor coupling through gamma (Ritterband-Rosenbaum et al., 2014). In addition, subjects that were led to believe they were producing sensory effects through their actions show modulations of gamma and beta oscillations associated with sensory processing (Buchholz et al., 2019). However, these previous studies have not manipulated the fluency of actions, which might be critical for agency (e.g. Sidarus 2017). In the automatic imitation task, we specifically manipulated the *congruency* between executed and perceived actions to boost or tamper actions. This manipulation has been effective for modulating the implicit feeling of agency (Vastano et al. 2017) and motor neural oscillations are likely play to reflect this effect.

In the current study we examined the dynamics of neural oscillations reflecting the congruency effects of automatic imitation on IB. We hypothesized a differential engagement of neural motor oscillations (alpha, beta and gamma) for actions as a function of congruency. In addition, correlations between neural and IB effects will indicate the relevance of these oscillations to implicit agency. Lastly, given the role of theta oscillations in inhibitory processes (e.g. Cohen and Ridderinkhof, 2013; Gulbinaite et al., 2014) and its potential involvement in automatic imitation task (Brass et al., 2000), we also explored modulation in theta oscillations. This secondary analysis will seek to better understand the neural processes involved in automatic imitation so far unexplored.

## 2. MATERIALS & METHODS

It should be noted that this study used the same dataset reported in our previous manuscript (Vastano et al., 2020). However, the nature of the data analysis differs substantially and none of the neural responses reported here were included in the previous study. In the current study we investigated the congruency effects of automatic imitation on neural oscillations rather than event-related potentials that were the focus of the previous study. For the sake of completeness we have included a brief description of the behavioural results (the reader can look in Vastano et al., 2020 for details).

### 2.1 PARTICIPANTS

Twenty-eight participants (age range: 18 to 28 years, mean 23.3 years, 20 female, 8 male) participated in the experiment after giving written informed consent. The study was approved by the local ethical committee of Ghent University and conducted according to the Declaration of Helsinki. All participants were right-handed and had normal or corrected-to-normal vision. All the participants were neurologically and psychiatrically healthy. Participants were paid for their participation.

### 2.2 STIMULI & TASK

The participants were seated in a comfortable armchair in a dim sound-attenuated room at a distance of 60 cm from the computer monitor (refresh rate 60 Hz, dimensions 53 × 30 cm and resolution 1920 × 1080). Visual stimuli were presented on a computer screen and auditory stimuli via headphones. Visual stimuli consisted of images (300 × 200 pixels) of a mirrored right-hand of an actor performing lifting finger movements (see Figure 1). At the start of a trial the message “Please, place your fingers” was displayed. The participants had to hold down the “G” and “H” keys of a Mac keyboard with numeric keypad (MB110Z/B) with their right index and middle fingers respectively. Once the participant placed her/his fingers the sequence continued. A fixation cross was presented between 1000 and 1600 ms and then an image showing a hand in a resting position was displayed for 1000 ms. This was followed by two simultaneous events: a number display (1 or 2 appearing between the two fingers) and the lift of one of the observed fingers (index or middle). Participants were instructed to respond as fast as possible to the ‘1’ by lifting the index finger and to the ‘2’ by lifting the middle finger. If no response was given within a time window of 1400 ms the next trial was presented and the missed trial was recovered. Following the key release and a variable interval (300, 400, or 500 ms) an auditory stimulus (a pure tone at 1000 Hz) was delivered for 300 ms by means of headphones. Next, after a variable interval between 300 and 800 ms a Visual Analogue Scale (VAS) appeared. The VAS ranged between 100 and 900 ms and has marks of 200 ms-intervals. In order to measure IB, participants were asked to estimate the interval of time between their actions (key release) and the ensuing tone using the VAS. For this time estimation task participants used the mouse to point in the VAS. They had a maximum of 5000 ms to give the answer. Finally, after a variable inter-trial interval (1000, 2000 or 3000 ms) the next trial started. The critical manipulation in this experiment is that the observed finger movements could result in a match or in a mismatch with the instructed finger movement. In congruent (C) trials the participant moves the index finger and sees an index finger movement (and similarly for middle fingers). In incongruent (IC) trials the participant moves the index finger and sees a middle finger movement (or alternatively moves the middle finger and sees an index finger movement). In addition, there was also a condition where the number was displayed but the fingers didn’t move (baseline; B). The experiment consisted of 240 randomized trials: 80 trials for each condition (congruent, incongruent and baseline), each of which was composed by 26-28 trials for each interval (300, 400, and 500 ms). The experiment was divided in 4 smaller blocks of 60 trials each (20 trials in each congruent, incongruent and baseline) to allow participants to rest between blocks. Before the main experiment the participants were trained for the time estimation task and then were familiarized with the task. For the training, participants listened two tones separated by 100 or 900 ms and then they were asked to indicate if the time elapsed between the two tones were 100 or 900 ms. For this training they received a feedback (correct or incorrect response). This short training phase (21 trials) aimed at capacitating the participants to discriminate between 100 or 900 ms. For the sake of the manipulation in the main experiment the participants were told the interval between their action and the tone was always random between 100 ms and 900 ms. Following the training, the participants were familiarized with 30 randomized trials (15 for each interval) identical to the main experiment. Once the instructions were clear the main experiment starts. The task was implemented in E-prime 2.0 Professional software (Psychology Software Tools, Pittsburgh, PA). The duration of the whole experiment was about 80 min.

**Figure 1:**
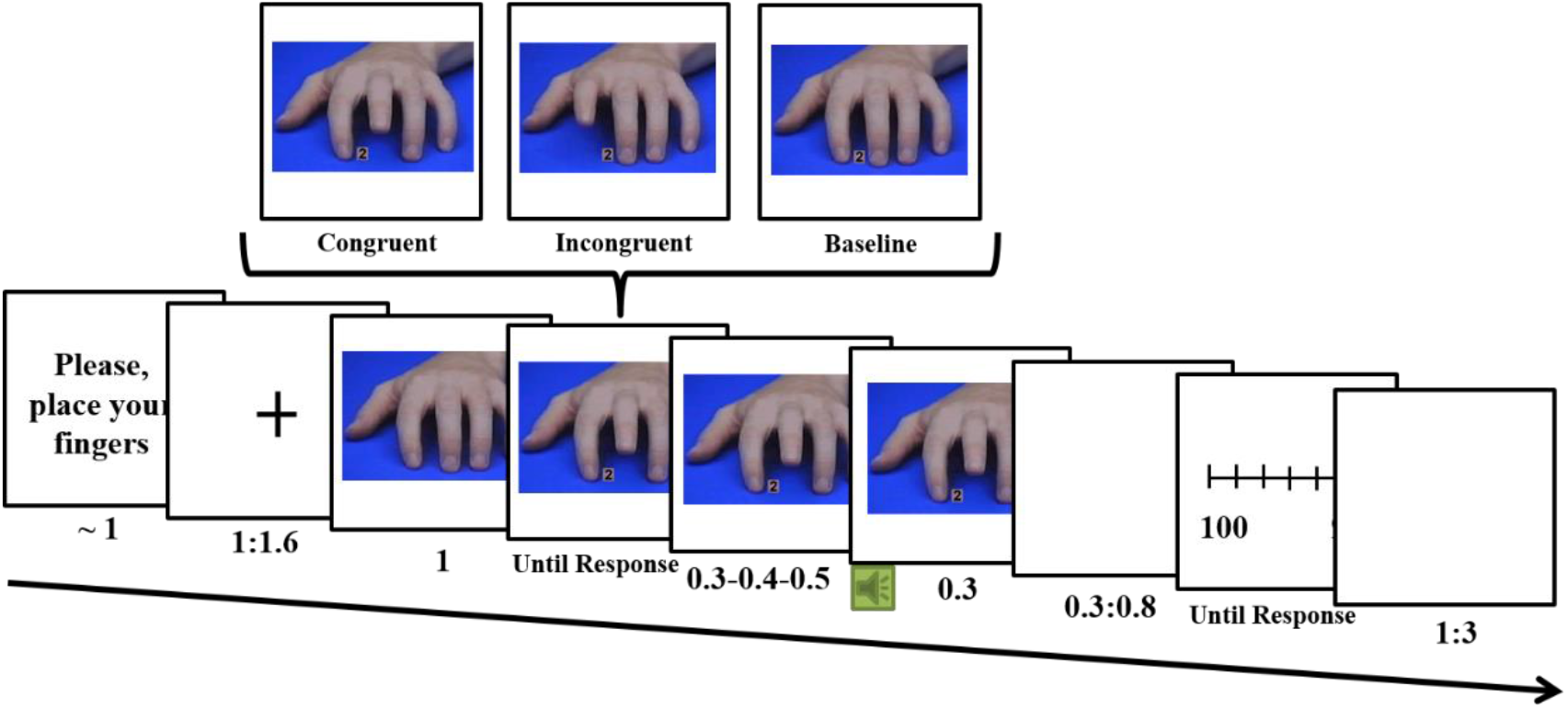
Time line for a trial. After placing their fingers and fixating a cross participants responded to a numeric cue displayed between the index and middle finger of a hand (this cue indicated which action participants should do). At the same time the displayed hand moved a finger that could be *Congruent* or *Incongruent* with the participant’ action. A condition where there was no movement of the displayed finger was also included (*Baseline*). After the participant’s response and a random interval (either 0.3, 0.4 or 0.5 s), followed a tone (the action outcome, the speaker in the figure). After a delay (between 0.3-0.6 s) a Visual Analogue Scale (VAS) ranging between 100 and 900 appeared. The participants had to judge on this VAS the time elapsed between their actions and the tone. Finally, after a variable inter-trial interval (between 1-3 s) the next trial started.

### 2.3 EEG RECORDING & SIGNAL PROCESSING

Scalp EEG was recorded from 64 active Ag-AgCl electrodes (Biosemi Active-Two, Biosemi, Amsterdam) mounted in an elastic cap according to the international 10–20 setting. The continuous EEG was recorded with a 1024 Hz sampling rate and referenced online to the CSM-DRL ground. Data was recorded in an electrically shielded chamber and electrode offsets were kept between −25 and 25 μV for all electrodes. Additional electrodes in a bipolar montage were applied near the canthi, and above and below the left eye to record the electrooculogram (EOG). EEG data analysis was performed using EEGlab 14.0.0b (Delorme and Makeig, 2004) and Fieldtrip (version 20170503; (Oostenveld et al., 2011) in MATLAB R2014b (The Mathworks, Inc, Natick, MA). The raw EEG data was loaded in EEGlab and filtered off-line.

#### 2.3.1 ARTEFECT REJECTION

In a first step raw data was filtered between 0.5 and 40 Hz. Non-stereotyped artefacts were cleaned and bad channels were detected by visual inspection. Bad channels (F6, FC6, T7, P2, and Iz across eight different participants) were interpolated using spherical splines (Perrin et al., 1989). Next, stereotyped artefacts (such as eye movements, eye blinks and muscle tension) were reduced by independent component analysis (ICA) and the SemiAutomatic Selection of Independent Components (SASICA) toolbox (Chaumon et al., 2015), removing no more than 3 components. ICA decompositions were performed separately on each subject over all conditions and then saved. In a parallel step, raw data was not filtered. The same corrections regarding non-stereotyped artefacts and bad electrodes done were re-applied to this dataset. Following this, the ICA weights computed before were also re-applied to this dataset. Applying the pre-computed weights allowed to efficiently remove artefacts while retaining the high-frequency EEG information (Artoni et al., 2017; Oddo et al., 2016; Winkler et al., 2015). Finally, this dataset was re-referenced to the average of all electrodes and visually checked one last time for artefacts. Any portion of data with remnant artefact was eliminated. The final dataset retained in average 66.5 trials (83%) per condition from the original raw data.

### 2.4 EEG ANALYSES

Time-frequency representations of power were estimated in single-trial data in two ways. For frequencies below 40 Hz we used short-time Fourier transform with sliding windows of 500 ms multiplied with a Hanning taper and moving steps of 50 ms. The frequency resolution was 2 Hz. For frequencies above 40 Hz we used a multitaper approach with a window length of 400 ms and frequency smoothing of 20 Hz (i.e. 15 tapers). Power was estimated as the square of the analytic signal z (power = real[z(t)]^2^ + imag[z(t)]^2^) and averaged across trials. Power values at each time-frequency point were normalized by converting to relative change ([Power_task-Power_baseline]/Power_baseline)) to account for power-law scaling of oscillations in different frequency bands. Time-frequency analyses were time-locked to the onset of the subject responses with a −1300 to 1400 ms time-down. Power from −750 to −1000 ms in the pre-stimulus period (when the observed hand stay still) served as the frequency band-specific baseline.

We averaged the data across all conditions and then created topographical plots for power in the alpha, beta, gamma and theta frequency bands in a 1-s window of interest, time-locked to the subject response (−0.5 s to 0.5 s). Second, electrodes that showed the largest change in the group-averaged power were selected. Third, condition-averaged and electrode-averaged time-frequency power plots were constructed for the selected electrodes for each frequency band. Fourth, time-frequency windows with the largest power increase were selected based on visual inspection (marked in Figures 2–5 as dashed squares in time-frequency plots). This selection procedure is independent of any condition-specific differences in power, and therefore could not have introduced any biases into the results (Gulbinaite et al., 2014). We also explicitly avoided to interpret any power changes at the edges of the scalp to minimize confounds related to any residual muscular activity (Goncharova et al., 2003). Finally, for each subject, the condition-specific power in the selected time windows of interest were used for statistical analyses.

### 2.5 STATISTICAL ANALYSES

Behavioural analyses have been performed previously in Vastano et al. (2020). For the EEG data we performed a repeated measures ANOVA on power estimates for each frequency with Congruency (3 levels: congruent, incongruent and baseline) as factor. All significant effects found in the ANOVAs were followed by Newman-Keuls corrected post hoc tests. Alpha level was fixed at 0.05 for all statistical tests. Greenhouse–Geisser correction was used to correct for sphericity violations when necessary.

## 3. RESULTS

In the automatic imitation task participants responded to a numerical cue that appeared between two fingers while one finger simultaneously moved (Figure 1). Following the response of the participants and after a delay the participants listened to a tone and estimated the time elapsed between their actions and the tone (this time estimation task allows us to measure IB). We set out to characterize the modulation of neural oscillations occurring during automatic imitation to investigate its impact on IB. Before presenting these results we will briefly present the behavioural results.

### 3.1 BEHAVIOURAL RESULTS

#### Performance and intentional binding increased for congruent actions

Behavioural results from this study have been reported in a previous publication (Vastano et al., 2020). Accuracy was higher for the congruent condition (98.4%) followed by the baseline (97.1%) and the incongruent (87.5%) condition. Similarly, reaction times were the fastest for the congruent condition (427 ± 63 ms, mean ± standard deviation) followed by the baseline (475 ± 70 ms) and incongruent (493 ± 78 ms) conditions. Thus, as typically observed in the automatic imitation task, performance of the participants is facilitated by congruent relative to the baseline and incongruent conditions.

In this task we evaluated whether time compression between actions and outcomes (the IB effect) would be modulated by congruency. Participants performed a time estimation task following their responses to the numeric cues. IB was measured as the difference between the participants’ estimations and the real interval durations (time judgements errors). Larger temporal compression reflects an increased IB effect. We observed that the congruent condition (−72 ± 81 ms) led to significantly reduced time judgment errors than the incongruent condition (−62 ± 74 ms), while a marginally significant difference was observed between the congruent and the baseline (−61 ms ± 80 ms) conditions. This means that congruency in the automatic imitation task can modulate IB levels. There is increased IB when the participants perform congruent compared to incongruent actions. Interestingly, the IB levels produced by incongruent and baseline trials is very similar, suggesting that facilitated actions can increase IB levels.

### 3.2 EEG RESULTS

#### Decreased gamma power for congruent actions

We inspected mid-central and left- and right-central electrodes where previous studies have found gamma activity (e.g. (Ball et al., 2008; Grent-’t-Jong et al., 2013; Muthukumaraswamy, 2010). We determined in which time-frequency windows the gamma power changes occurred across all conditions. We observed a peak of gamma power confined at electrodes FC1, FCz, FC2, C1, Cz and C2 in a time-frequency range between 70 and 95 Hz and −0.2 and 0.2 s relative to response onset (see Figure 2A and 2B, “All” conditions). We then examined whether gamma activity would be modulated by congruency. We found a significant effect of Congruency (F(2,54) = 3.57; p = 0.035; η^2^p = 0.21) with decreased gamma power for the congruent (C; Mean = 0.009, SD = 0.033) relative to the incongruent condition (IC; Mean = 0.032, SD = 0.031; p = 0.039). Gamma power for the incongruent condition was not different from the baseline condition (B; Mean = 0.024, SD = 0.033; p = 0.38; Figure 2). Executed actions that match the observed actions led to decreased gamma activity relative to the incongruent condition, suggesting that gamma activity differentiates between actions that were congruent or not with observed actions.

**Figure 2:**
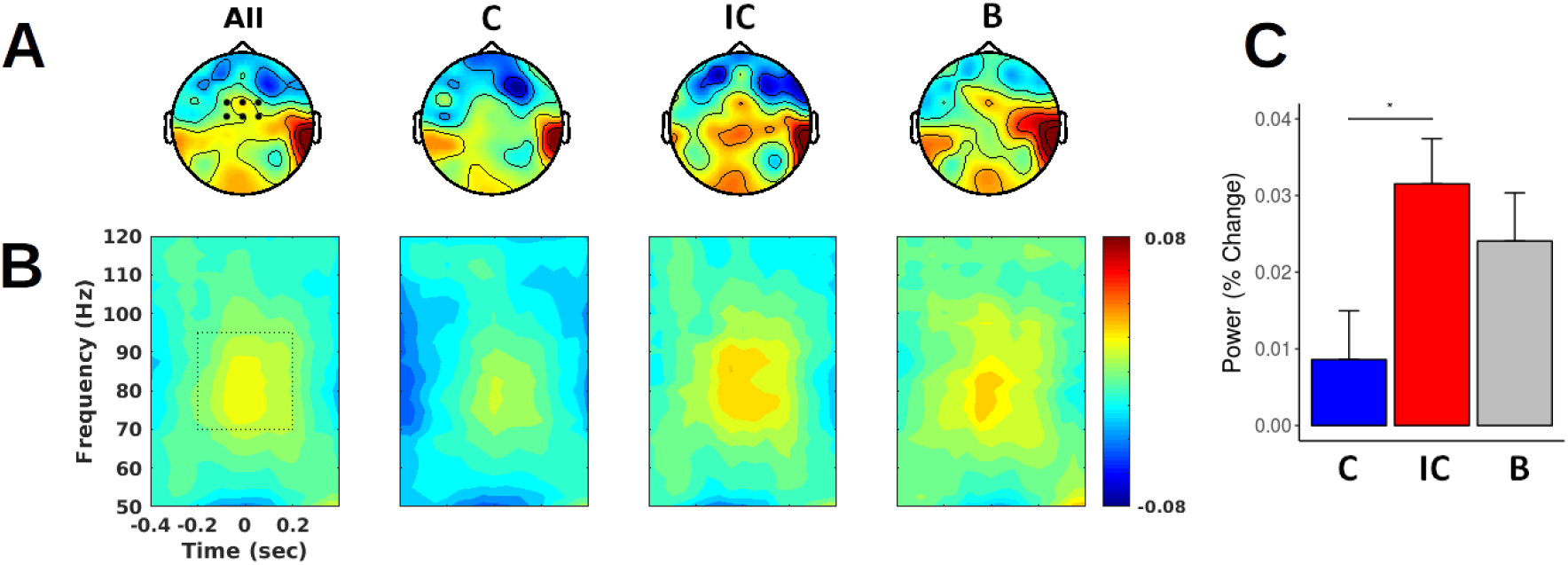
Gamma power is less for congruent compared to incongruent actions. A) Topographic distributions of grand-average gamma (70-95 Hz) activity in the −0.2 and 0.2 s interval relative to response onset for all (All), congruent (C), incongruent (IC), and baseline (B) conditions. B) Time-frequency representation of power estimates at electrodes depicted in A. C) Bar plot of grand-average gamma activity for congruent (blue), incongruent (red) and baseline (grey) conditions. Error bars represents standard errors of the mean. * = p < .05.

#### Gamma power differences correlates with intentional binding effects

Our second aim was to investigate whether gamma power changes were related to the IB effects. To assess this relationship we subtracted for each subject the mean power of congruent trials from the mean power of incongruent trials (Power_IC_ – Power_C_). Similarly we computed an IB effect by subtracting for each subject the mean IB value of congruent trials from the mean IB value of incongruent trials (IB_IC_ – IB_C_). We used Spearman correlations to investigate the relationship between gamma power differences with IB effects over participants. The gamma power differences correlated positively with the IB effect (r(26) = 0.47; p = 0.012; Figure 3). Acknowledging that correlational analyses can be sensitive to deviant observations and characteristics of the dataset and that these analyses can be flawed by the presence of outliers (Rousselet and Pernet, 2012) we overcome this problem by conducting a robust regression analyses using the Robust Correlation toolbox (Pernet et al., 2013). Within this toolbox the skipped correlation analysis takes the overall structure of the data into account protecting against the detrimental effects of aberrant observations while maintaining the false positive rate below the nominal alpha level (Pernet et al., 2013; Wilcox, 2004). The gamma power differences still correlated positively with the IB effect (r = 0.5; h = 1). This result indicates that those participants that show greater congruency effects in IB (greater time compression for congruent compared to incongruent actions) also presented greater gamma power decreases for congruent versus incongruent actions.

**Figure 3:**
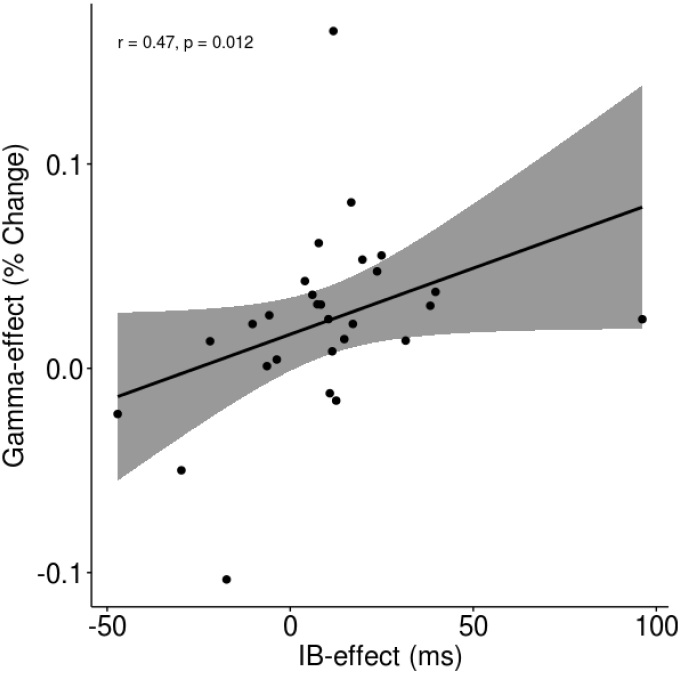
Gamma power differences correlated positively with the intentional binding effects. Dispersion plot and tendency line for the gamma power effect versus the IB effect (IB_IC_ – IB_C_). Each dot represents a participant.

Next, we implemented a post-hoc exploratory analysis to determine whether the relation between gamma power and IB was dominated by a facilitatory and/or an interference effect. For the neural facilitatory effect we subtracted for each subject the mean power of congruent trials from the mean power of baseline trials (Power_B_ – Power_C_). Similarly we computed a behavioural facilitatory effect by subtracting for each subject the mean IB value of congruent trials from the mean IB value of baseline trials (IB_B_ – IB_C_). We then correlated the neural facilitatory effects with the behavioural facilitatory effects over participants. To investigate an interference effect we compute similar neural and behavioural differences, but subtracting baseline trials from incongruent trials (IC – B). The facilitatory effect for gamma power correlated positively with the behavioural facilitatory effect (r(26) = 0.44; p = 0.02), but no relationship was found between the interference effect for gamma power and the behavioural interference effect (r(26) = 0.15; p = 0.44). This suggests that the link between gamma power and IB effects might be driven by a facilitatory effect. However, conducting a robust correlation these effects become non-significant (r = 0.43; h = 0). Taking altogether, we observe an overall effect where those participants that show greater time compression for congruent versus incongruent actions also presented greater gamma power decreases. We also observe a trend where participants that showed greater increased facilitatory effects in IB also showed greater gamma power decreases. This trend is in favour of the notion that changes in gamma power on IB might be driven by a facilitatory effect albeit the current data do not allow us to make a firm conclusion.

#### Decreased alpha and beta power for congruent actions

Next, we characterized changes in alpha and beta power in the automatic imitation task. We determined the time-frequency windows where changes in alpha and beta occurred across all conditions. We observed a peak of alpha power confined at electrodes CP3, CP1, P3, P1 and C3 (left-central side) and CP4, CP2, P4 and C4 (right-central side) in the 8 to 12 Hz band and between −0.2 and 0.2 s relative to response onset (Figures 4A and 4B, “All” conditions). We then examined whether alpha activity would be modulated by congruency. Given that the participant responded with the right hand we focussed our analyses on the left hemisphere. We found a significant effect of Congruency (F(2,54) = 5.03; p < 0.01; η^2^p = 0.29) with lower alpha power for the congruent (C; Mean = −0.19, SD = 0.24) relative to the incongruent (IC; Mean = −0.12, SD = 0.32; p = 0.013) and baseline condition (B; Mean = −0.14, SD = 0.31; p = 0.037; Figure 4). This suggests that the alpha activity might reflect the similarity between motor representations evoked by executed and observed actions in the automatic imitation task.

**Figure 4:**
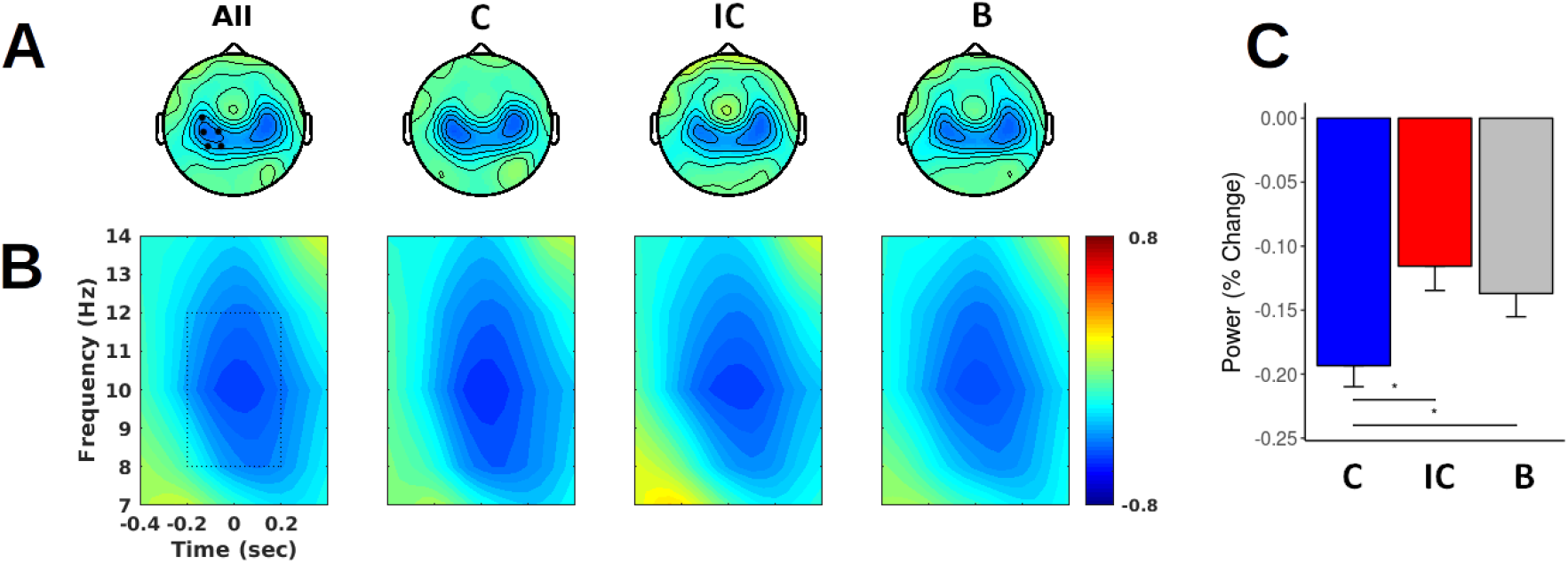
Alpha power is lower for congruent actions. A) Topographic distributions of grand-average alpha (8-12 Hz) activity between −0.2 and 0.2 s relative to response onset for all (All), congruent (C), incongruent (IC), and baseline (B) conditions. B) Time-frequency representation of power estimates at electrodes depicted in A. C) Bar plot of grand-average alpha activity for congruent (blue), incongruent (red) and baseline (grey) conditions. Error bars represents standard errors of the mean. * = p < .05.

We observed that a peak of beta power confined at electrodes C3, C1, CP3 and CP1 (left-central side) and C4, C2, CP4 and CP2 (right-central side) in the 15 to 30 Hz beta band between −0.3 and 0.2 s relative to response onset (Figure 5A and 5B, “All” conditions). Next, we examined whether beta activity would be modulated by congruency. Given that participants respond with the right hand we focussed the analyses on the left hemisphere. We found a significant effect of Congruency (F(2,54) = 12.04; p < 0.001; η^2^p = 0.49) with reduced beta power for the congruent (C; Mean = −0.29, SD = 0.13) relative to the incongruent (IC; Mean = −0.24, SD = 0.14; p < 0.001) and baseline conditions (B; Mean = −0.24, SD = 0.14; p < 0.001; Figure 5). This suggests that decreased beta activity, in a similar manner as alpha activity, might signal the match between performed and executed motor representations in the automatic imitation task.

**Figure 5:**
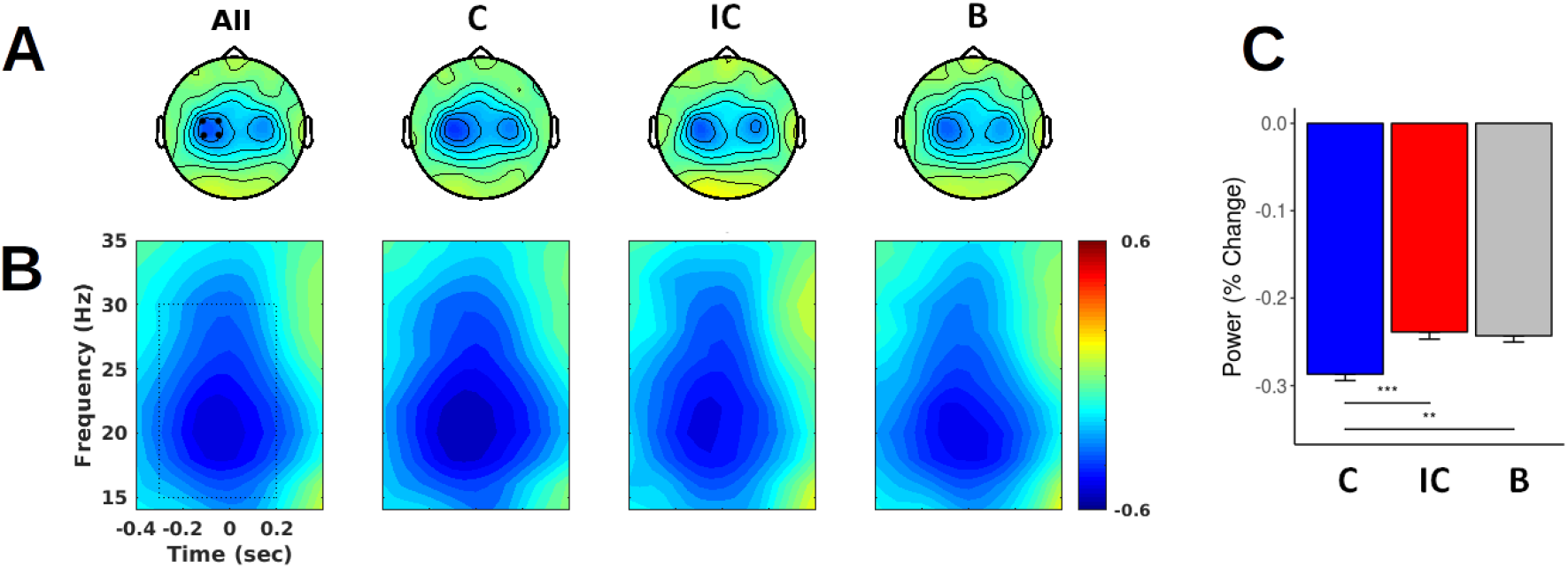
Beta power is lower for congruent actions. A) Topographic distributions of grand-average beta (8-12 Hz) activity between −0.3 and 0.2 s relative to response onset for all (All), congruent (C), incongruent (IC), and baseline (B) conditions. B) Time-frequency representation of power estimates at electrodes depicted in A. C) Bar plot of grand-average beta activity for congruent (blue), incongruent (red) and baseline (grey) conditions. Error bars represents standard errors of the mean. ** = p < 0.01; *** = p < 0.001.

Following this characterization, we investigated whether alpha and beta power changes were relevant for IB effects. To assess this relationship, we performed the same correlations analyses described for gamma power. We correlated differences in alpha and beta power with IB effects over participants. Alpha and beta power differences did not correlated with the IB effects (alpha: r(26) = −0.03; p = 0.86; beta: r(26) = −0.29; p = 0.14).

#### Increased theta power for incongruent actions

Lastly, we characterized changes in theta power in the automatic imitation task. We determined the time-frequency windows where changes in theta activity occurred across all conditions. We observed a peak of theta power confined at FCz in the 4 and 8 Hz band between −0.2 and 0.1 s relative to response onset (see Figure 6B, “All” conditions). Next, we examined whether theta power was modulated by congruency. We found a significant effect of Congruency (F(2,54) = 43.15; p < 0.001; η^2^p = 0.75), with a strong increase in theta power for the incongruent (IC; Mean = 0.98, SD = 0.58) relative to the congruent (C; Mean = 0.46, SD = 0.39; p < 0.001) and baseline (B; Mean = 0.57, SD = 0.5; p < 0.001; Figure 6) conditions. These findings are in agreement with previous research that indicates that theta oscillations might be involved in conflict processing, as depicted in the increment of theta in the incongruent condition.

**Figure 6:**
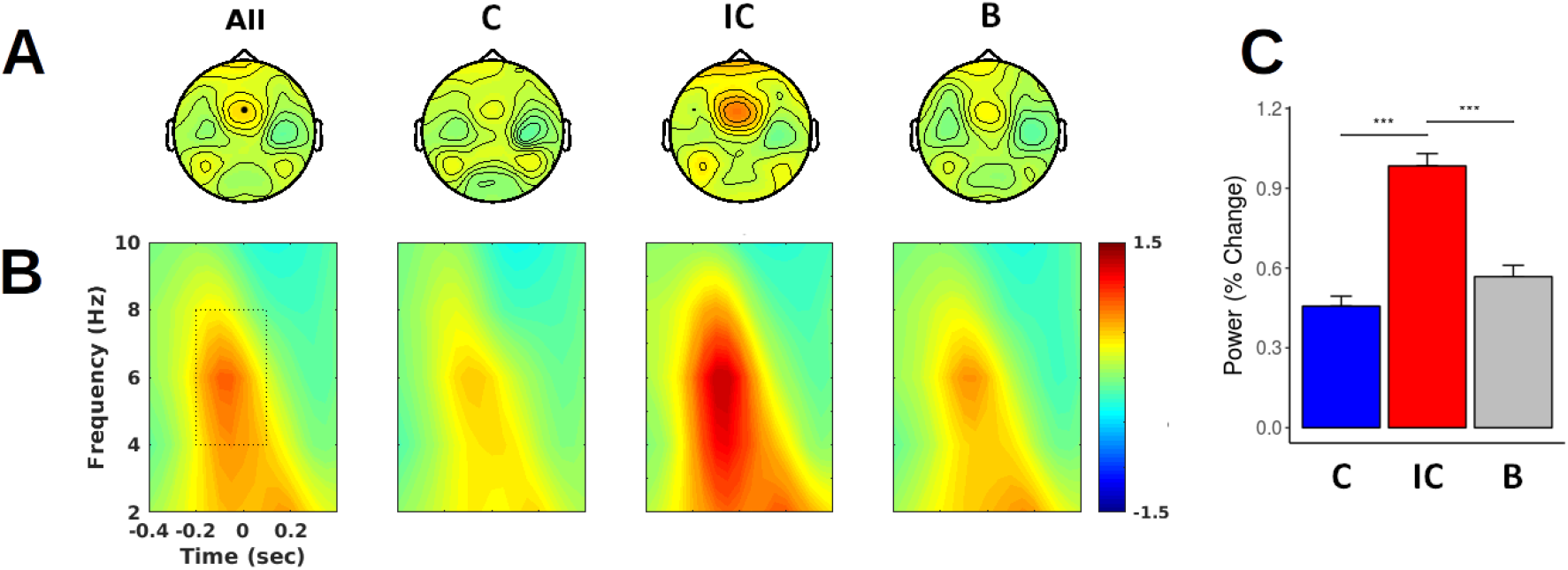
Theta activity. A) Topographic distributions of grand-average theta (4-8 Hz) activity between −0.2 and 0.2 seconds relative to response onset for all (All), congruent (C), incongruent (IC), and baseline (B) conditions. B) Time-frequency representation of power estimates at the FCz electrode (black dot in A) for the same conditions as in A. C) Bar plot of grand-average theta activity for congruent (blue), incongruent (red) and baseline (gray) conditions. Error bars represents standard errors of the mean. *** = p < 0.001.

Following this characterization, we investigated whether theta power changes were relevant for IB effects. To assess this relationship, we performed the same correlations analyses described for gamma power. We correlated differences in theta power with IB effects over participants. Theta power differences did not correlated with the IB effects (r(26) = 0.21; p = 0.27). Altogether, alpha and beta power together with theta power show no reliable relationship to IB.

## 4. DISCUSSION

We aimed at investigating the role of neural oscillations generated in sensory-motor regions in relation to the implicit feeling of agency as well as how they are modulated by automatic imitation. We manipulated the congruency between observed and planned actions and quantified the temporal compression (between actions and tones) experienced by the participants (the intentional binding, IB, effect). Participants have stronger IB when they performed congruent rather than incongruent actions in relation to the observed finger movement. Furthermore, participants that showed greater IB for congruent versus incongruent actions (congruency effect) also presented greater gamma power differences for congruent versus incongruent actions. Increased power desynchronization in alpha and beta was observed for congruent relative to incongruent trials, while increased power synchronization was observed in theta for incongruent relative to congruent trials. These frequencies were modulated by congruency but were unrelated to IB. In the sections below we will discuss the putative role of gamma oscillations for implicit agency and the significance of our findings for understanding automatic imitation.

### Implicit agency is related to gamma power changes

Our primary finding is that the implicit agency (as measured by IB) was associated with changes in gamma oscillations. Previous studies have shown the involvement of neural oscillations in agency (Kang et al., 2015; Ritterband-Rosenbaum et al., 2014), but these studies showed mostly sensory effects possible related to manipulations of sensory feedback. In our study, we manipulated the congruency of executed and observed finger movements, thus modifying the fluency of response selection. We expected to find modulations across our three candidate neural oscillations (alpha, beta and gamma). However, we only found that modulations in gamma power were associated with IB. The changes in alpha and beta power were unrelated to IB and seem to reflect mirror activity (see below). Previous studies suggest that the motor system is involved in the mental representations of actions performed by oneself and by others. However, by its very nature these resonance mechanisms cannot realise the function of self/other distinction (Brass and Heyes, 2005) which may be necessary for processes like agency. Thus, a possible interpretation is that mirror activity does not contribute to agency in this task. This aspect needs to be investigated further. The effect of *congruency* on gamma oscillations was associated with IB. Participants with greater congruency effects in IB (i.e. greater time compression differences between congruent and incongruent actions) also presented greater differences of gamma power between congruent and incongruent actions. One interpretation is that gamma oscillations may have a role in agency possibly reflecting monitoring of actions. In the motor system gamma oscillations show transient activities tightly locked to movement onset (Ball et al., 2008; Cheyne, 2013; Cheyne et al., 2008; Crone et al., 1998b) and have been thought to reflect the initialization of motor actions (Cheyne and Ferrari, 2013). Gamma oscillations also signal the competition of distinct motor programs in tasks where alternative responses are required (Gaetz et al., 2013; Grent-’t-Jong et al., 2013; Heinrichs-Graham et al., 2018; Isabella et al., 2015). This emergent evidence suggests that gamma oscillations might be critical for cognitive control in the motor domain. Crucially, facilitating or interrupting actions has been seen to affect agency. In previous priming studies an increment of fluency of actions resulted in increased judgements of agency (Chambon and Haggard, 2012; Chambon et al., 2014; Sidarus et al., 2017). Moreover, interrupting the normal flow of action processing with transcranial magnetic stimulation over pre-SMA or the parietal cortex decreases agency (Chambon et al., 2014; Zapparoli et al., 2020). Our findings suggest that gamma power may be able to track the effects of *congruency* (between perceived and performed actions) affecting the processing of actions and then its impact on implicit agency. A facilitation interpretation is suggested by the correlation (although non-significant) of the effect of congruent actions against the baseline condition. In addition, a previous study on automatic imitation has shown a decrease in the readiness potential for the congruent action (relative to the other conditions), supporting the notion that congruency effects are driven by action facilitation (Deschrijver et al., 2017). It is also worth noticing that in our previous study the correlation between the IB effect and late P300 changes was also driven by a facilitatory effect (Vastano et al., 2020). Increased late P300 amplitudes for congruent actions were interpreted as reflecting reduced cognitive load and accessible attentional resources. Together our ERP and gamma band neural oscillations findings suggest that monitoring and executive attention processes (both building blocks of executive functions) are affected by the congruency between observed and executed actions and these processes in turn affect implicit agency.

One distinctive aspect of this study is worth mentioning. There is a strong theta response elicited for the incongruent relative to the congruent condition. Theta responses have been typically associated with cognitive control and executive processes including response inhibition (Cohen and Ridderinkhof, 2013; Gulbinaite et al., 2014; Nigbur et al., 2011). Although here we focused neural oscillations from sensorimotor regions, theta oscillations could have also contributed to implicit agency insofar these are neural responses that signal interference or conflict and could affect fluency (Cohen, 2014). We correlated the congruency effects on IB with the theta effects but we did not find any relationship. This is surprising given that gamma oscillations correlate with IB and we interpret that these effects are associated with action facilitation. However, previous evidence shows that theta and gamma oscillations might reflect distinct aspects of cognitive control. Isabella et al. (2015) used a modified go/no-go task that included a switching condition and shows that theta oscillations were broadly related to cognitive control (rather than to response inhibition or switching) while gamma oscillations were specifically associated with switching responses. The increase of gamma power for the switch condition could be analogous to the incongruent condition in our automatic imitation task. This again highlights the role that gamma oscillations can have for monitoring and updating motor programs, in complement to theta oscillations. Future studies will need to further investigate this interesting aspect of motor control. The role of gamma oscillations should also be related to its location in the cortex. Gamma oscillations have been shown to be generated in primary motor cortices but also in brain areas such as the supplementary motor area (SMA) and the premotor cortex (Ball et al., 2008; Szurhaj et al., 2005). In our study, gamma effects seem to be produced by central generators in the brain (although EEG does not allow to draw strong conclusions). Interestingly, the SMA has an important role in voluntary movements (Deiber et al., 1991, 1996) and timing perception (Wiener et al., 2010). In fact, activity in the SMA is directly related to the time compression between voluntary actions and outcomes experienced by participants in a time estimation task (Kühn et al., 2013). In sum, our study suggests that as we perform actions we have an implicit feeling of agency that is sensitive to the fluency experienced in these actions. Our brain is able to detect these changes through the implementation of monitoring processes involving fast rhythms in the gamma band range.

### Neural oscillations in automatic imitation

Our secondary finding is related to the modulation of neural oscillations by congruency in automatic imitation. Importantly, this study is the first that investigates oscillatory pattern in this task. During this task a participant responded to a visual cue while observing a finger movement produced by a mirrored hand in the background and where the participant’s response could be congruent or incongruent with the observed finger movement (or “neutral” if no movement was observed). We identified strong movement-locked responses in the alpha, beta and theta bands. Alpha oscillations at central brain areas have been typically associated with perception of biological movement (Perry & Bentin 2009) and beta oscillations have been related to motor preparation (Pfurtscheller et al., 1997), amongst other functions. In our task, alpha and beta oscillations show increased desynchronization for the congruent relative to the incongruent condition, possibly reflecting increased visuo-motor integration and processing for movements that match (relative to movements that do not match) the observed actions. This effect could reflect the engagement of a mirror neuron system (MNS), neurons in premotor areas that are more active both when an individual observes and executes similar actions (di Pellegrino et al., 1992; Perry and Bentin, 2009; Pineda and Hecht, 2009). This is consistent with previous fMRI (Cross et al., 2013; Iacoboni, 1999) and MEG (Kessler et al., 2006) studies showing the involvement of the MNS in automatic imitation. The involvement of both alpha and beta oscillations may imply mirror activities in these two frequency bands (Muthukumaraswamy and Singh, 2008; Rossi et al., 2002). Finally, in our study, theta oscillations possibly signal the increased requirement for inhibition of imitative tendencies in the incongruent condition. This is consistent with previous studies on automatic imitation showing the involvement of frontal and central brain areas associated with conflict processing (Bien et al., 2009; Brass et al., 2001, 2005; Cross et al., 2013, see also Darda and Ramsey, 2019). Altogether, our findings contribute to a better understanding of the neural underpinnings of automatic imitation. Future studies should aim at better characterize interactions between the distinct rhythmic activities that occur in automatic imitation, for instance between gamma and theta oscillations (Jensen and Colgin, 2007).

We must acknowledge some limitations of our study. First, unlike previous work on agency we focused on evaluating implicit agency using a time estimation task. The relationship between the implicit feeling and the judgement of agency is still not fully clarified (Synofzik et al., 2008) but see (Ebert and Wegner, 2010) so we cannot make strong conclusions regarding the general mechanisms of agency. A fruitful avenue of research when investigating the neural correlates of agency will be to include these two types of agency. Second, since we focused on neural responses time-locked to the motor responses it could argued that differences of neural activities between conditions are merely related to differences in motor parameters. We think this is unlikely given that all conditions evoked a similar uplift finger movement and the difference only concern the congruency between observed and executed actions. Future studies should include additional measures of movements such as electromyography to better monitor these effects. Lastly, since there is evidence that agency is impaired in psychopathologies such as schizophrenia (Frith, 2005; Voss et al., 2010) it will be interesting to investigate neural changes associated with this task in psychopathologic populations.

## CONCLUSION

We used an automatic imitation task and EEG to investigate oscillatory responses associated with the implicit feeling of agency. Modulations of IB were associated with gamma power changes, suggesting a role of gamma oscillations in the implicit feeling of agency. In addition, during automatic imitation increased sensorimotor integration was reflected in alpha and beta oscillations, while general conflict mechanisms were reflected in theta oscillations. These findings provide novel insight into how neural oscillations are involved in automatic imitation.

